# Epigenetic modulation of RIPK3 by Transglutaminase 2-dependent serotonylation of H3K4me3 affects necroptosis

**DOI:** 10.1101/2024.10.18.619049

**Authors:** Alessio Vecchio, Fiorella Colasuonno, Veronica Bellanca, Consuelo Pitolli, Luca Occhigrossi, Fabio Ciccarone, Manuela D’Eletto, Federica Di Sano, Vittoria Pagliarini, Claudio Sette, Mauro Piacentini, Federica Rossin

## Abstract

The receptor interacting protein kinase 3 (RIPK 3) is the main player in the activation of necroptosis, a pro-inflammatory regulated cell death modality induced by many different stimuli. RIPK3 is epigenetically regulated by DNA methylation and can be expressed when its promoter is associated with H3K4me3 histone. In this study, we show that Transglutaminase 2 protein (TG2) is necessary to induce necroptosis pathway allowing the expression of *Ripk3* gene. Indeed, cells lacking TG2 show a strong downregulation of *Ripk3* gene and are resistant to necroptotic stimuli. TG2 is known to promote the serotonylation of H3K4me3 histone (H3K4me3Q5ser) regulating in this way the target gene expression. Interestingly, we find that TG2 interacts with both histones H3K4me3 and H3K4me3Q5ser and these post-translational modifications are associated with the *Ripk3* promoter only in presence of TG2. In addition, the absence of the H3K4me3Q5ser, in cells lacking TG2, is correlated with *Ripk3* gene methylation. Altogether, these results indicate that RIPK3 expression requires TG2 mediated serotonylation of H3K4me3 to prevent *Ripk3* promoter methylation, thus favouring its expression and necroptosis induction.

**HIGHLIGHTS:**

- Transglutaminase 2 protein (TG2) is necessary to induce necroptosis.
- Cells lacking TG2 show a strong downregulation of *Ripk3* gene.
- TG2 allows Ripk3 gene expression by promoting the serotonylation on H3K4me3 histone.

**SYNOPSIS:** 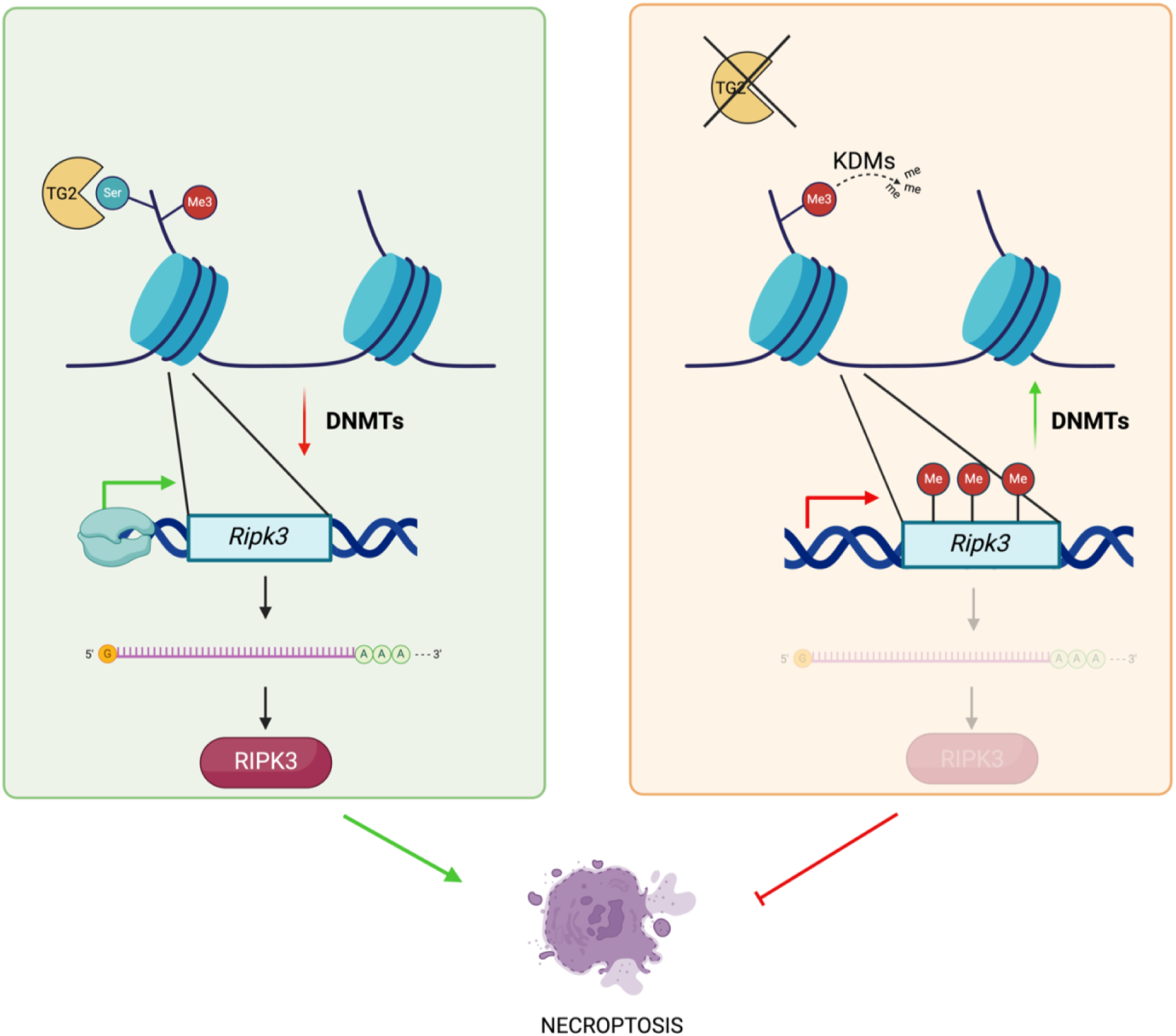

- TG2 serotonylates histone H3K4me3, associated to *Ripk3* promoter, favoring its expression.
- TG2 dependent serotonylation of histone H3K4me3 prevents DNA methylation.
- TG2 favours the expression of Ripk3 making cells susceptible to necroptosis.
- In the absence of TG2, neither serotonylation nor methylation occur on histone H3, associated to Ripk3 promoter.
- In the absence of TG2, Ripk3 promoter is methylated, thus blocking the activation of necroptosis.

## INTRODUCTION

Necroptosis is an inflammatory-mediated form of cell death that occurs with a necrotic morphotype because of the plasma membrane rupture, but this phenomenon is finely regulated at the molecular level, similar to apoptosis [1, 2]. Necroptosis can be triggered by various extracellular stimuli, known to activate inflammation and cell death, such us TNFα, LPS, and others. Inflammatory signals, through the binding of the TNFα to its receptor, promote the phosphorylation of Receptor Interacting Protein Kinases 1 (RIPK1), the interaction between RIPK1 and RIPK3, self-phosphorylation of RIPK3 which in turn phosphorylates the Mixed Lineage Kinase domain-Like protein (MLKL). Phosphorylated MLKL oligomerizes and translocates to the plasma membrane forming pores thus causing its rupture [3, 4].

RIPK3 is a key player in the necroptosis pathway and its expression can be regulated both transcriptionally, by the transcription factor Sp1, and epigenetically by histone modifications and DNA methylations, that are responsible for the expression and/or the repression of the gene [5–8]. In a recent study, performed in mouse embryonal fibroblasts (MEFs), it has been demonstrated that *Ripk3* gene can be expressed only when its promoter is associated with the trimethylation on lysine 4 of H3 histone (H3K4me3) [6]. Transglutaminase 2 (TG2) is a multifunctional enzyme implicated in several cellular processes such as cell growth, differentiation, autophagy and cell death [9–10]. High concentrations of Ca^2+^ and low concentrations of GTP are required for the ТG2 transamidating/crosslinking activity [11]. Moreover, TG2 has also calcium independent activities such as G-protein, scaffold, kinase, and protein disulphide isomerase activity [11–12]. Many groups have demonstrated that nuclear TG2 can catalyze the post-translational modifications of important transcription factors as p53, pRB, Sp1 and HSF1 thus playing a direct key regulatory role on gene transcription [13–14]. Despite the publication of the first compelling evidence of the nuclear presence of TG2 over four decades ago, little attention has been paid to its potential epigenetic regulatory function [14–15]. Different studies demonstrated that glutamines, present in the histones, act as amine acceptors in the transamidating reactions catalysed by TG2 [16–17]. However, only recently new molecular evidence has emerging about TG2 as a potential epigenetic regulator. In fact, it has been convincingly demonstrated that TG2 can add the serotonin molecule on glutamine 5 of H3K4me3, the histone 3 tri-methylated at the 4th lysine residue. This TG2-dependent post-translational modification (H3K4me3Q5ser) plays a key role in favouring gene expression [18]. Interestingly, just in the last months ripk3 has been listed among the genes whose promotor is associated with H3K4me3Q5ser modification, in a screening of placenta samples [19].

In this work, we demonstrate that TG2 is required for the activation of the necroptosis pathway defining a new molecular mechanism responsible for the *Ripk3* gene expression. Our results indicate that TG2, by promoting the serotonylation on H3K4me3 histone, allows *Ripk3* gene expression and consequently makes cells susceptible to necroptosis.

## RESULTS AND DISCUSSION

### The absence of TG2 leads to an impairment of the necroptotic pathway

We recently elucidated the overall impact of TG2 on gene expression by performing the RNA-seq analysis of WT MEFs and TG2 KO MEFs [10]. Unexpectedly, the results showed about 5000 genes that significantly changed their expression in TG2 KO MEFs. Interestingly, among them, we identified a strong downregulation of *Ripk3* mRNA levels in MEFs lacking TG2 (–192,6 fold) compared to WT MEFs [10]. Prompted by these findings, we first confirmed the reduced mRNA expression of *Ripk3*, by Real Time qPCR (RT-qPCR), in MEF cell lines. Effectively, we found a strong reduction of *Ripk3* mRNA levels in TG2 KO MEFs with the respect to WT MEFs (Figure S1A). Moreover, the analysis of the protein expression levels, by western blot, revealed the absence of RIPK3 protein in TG2 KO MEFs (Figure S1B), thus confirming that in absence of TG2 RIPK3 is not expressed.

RIPK3 is a main player in necroptosis pathway, thus we wanted to verify the activation of the pathway in MEFs lacking TG2 by analysing the phosphorylation of RIPK1, RIPK3 and MLKL, initial events in the necroptosis process. To induce necroptosis, we used tumour necrosis factor α (TNFα), SMAC mimetics (to inhibit the formation of complex I that promotes cell survival by NF-kB) and Z-VAD-FMK (to block complex IIa formation and thus apoptosis by caspase 8 inhibition), hereafter referred as “TSZ”. To this aim, we treated WT MEFs and TG2 KO MEFs with TSZ for 1, 3 and 6 h. After the TSZ treatment, we found the phosphorylation of RIPK1 (ser166) in both MEF cell lines (Figure 1A), but we did not observe the phosphorylation of RIPK3 and MLKL in TG2 KO MEFs (Figure 1B). As expected, the impairment of the phosphorylation of the key players of the necroptotic pathway, observed in MEFs lacking TG2, was also associated to a marked reduction in cell death (Figure S1 C). To verify that the decreased cell death was due to a block of necroptosis, we analysed the presence of the oligomeric form of MLKL, which is essential for the creation of membrane pores leading to plasma membrane rupture. The western blot analysis revealed the existence of MLKL oligomers exclusively in WT MEFs and not in TG2 KO MEFs (Figure 1C). To validate these findings, we performed an immunofluorescence analysis of MLKL and we observed an increase in the immunofluorescence intensity levels of MLKL in WT MEFs treated with TSZ for 3 h as compared to TG2 KO MEFs (Figure 1D). This increase was paralleled by the enhanced formation of green dots that are more pronounced in WT MEFs indicating the oligomerization of MLKL (Figure 1D, arrows). Collectively, these results suggest that in the absence of TG2, necroptosis may not be activated, due to the impaired expression of RIPK3.

**Figure 1.**
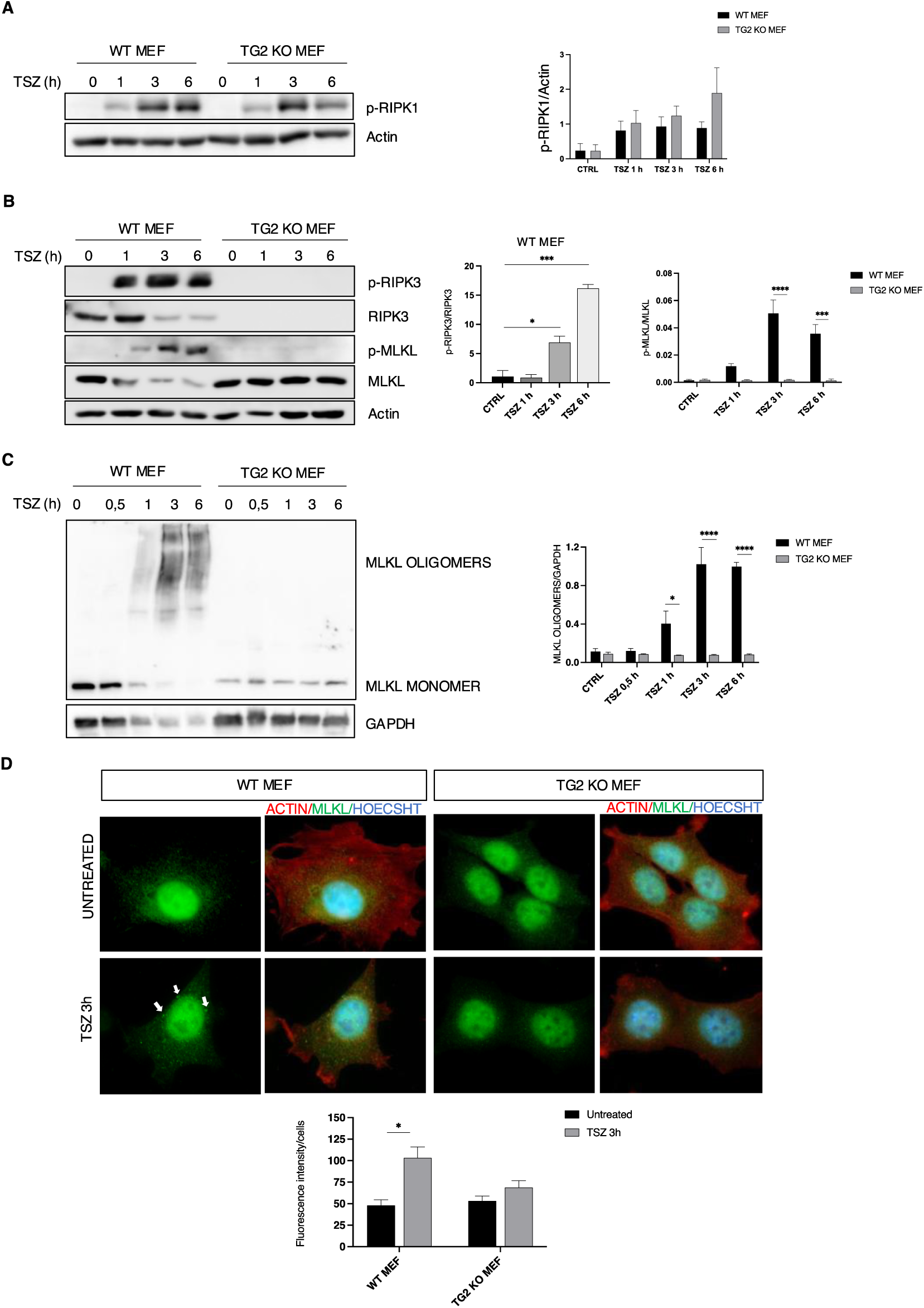
In absence of TG2 the induction of necroptosis is impaired. (A) Representative western blot and densitometric analysis of P-RIPK1 in WT and TG2 KO MEFs after TSZ treatment for 1, 3 and 6 h. Actin was used as loading control. The graph shows the mean ± SEM of densitometric analysis from three independent experiments. The P-value was determined by two-way ANOVA with Sidak’s multiple comparation test. (B) Representative western blot and densitometric analysis of P-RIPK3, RIPK3 and P-MLKL in WT and TG2 KO MEFs after TSZ stimulation for 1, 3 and 6 h to induce necroptosis. Actin was used as loading control. The graph shows the mean ± SEM of densitometric analysis from three independent experiments. The P-value was determined by unpaired Student’s *t* test and two-way ANOVA with Sidak’s multiple comparation test (*p < 0.05, ***p < 0.001, ****p < 0.0001). (C) Representative western blot and densitometric analysis of MLKL oligomers in WT and TG2 KO MEFs after TSZ treatment for 0.5, 1, 3 and 6 h. The protein samples were treated in a non-reducing condition. GAPDH was used as loading control. The graph shows the mean ± SEM of densitometric analysis from three independent experiments. The P-value was determined by two-way ANOVA with Sidak’s multiple comparation test (****p < 0.0001). (D) Immunofluorescence analysis performed after TSZ stimulation for 3 h, using antibodies against MLKL (green) and β actin (red). Nuclei are stained with Hoechst (blue). Representative images, captured at 60X magnification. Quantification of fluorescence intensity levels of MLKL analysed by ImageJ (NIH) software. The graph shows the mean ± SEM of fluorescence intensity from three independent experiments. The P-value was determined by two-way ANOVA with Sidak’s multiple comparation test (*p < 0.05).

### Silencing of TG2 in WT MEFs mitigates necroptosis

The previous results suggest that TG2, favouring RIPK3 expression, is required for necroptosis activation. To confirm this evidence, we silenced TG2 expression to further validate its effect on RIPK3 expression and necroptosis induction. To this aim, we ablated TG2 in WT MEFs by RNA interference for 72 h in basal conditions. Interestingly, we observed a decrease of RIPK3 protein levels in MEFs silenced for TG2 (siTG2) compared to the scramble controls (scr) (Figure 2A). We also verified whether the decreased TG2 expression could mitigate necroptosis. As expected, after TSZ treatment for 1, 3 and 6 h we observed a decreased phosphorylation of RIPK3 (Figure 2B) and MLKL (Figure 2C) in MEFs silenced for TG2 with the respect to the scramble conditions together with a decrease of the oligomeric form of MLKL (Figure 2D). These results confirm that TG2 promotes necroptosis by modulating RIPK3 expression.

**Figure 2.**
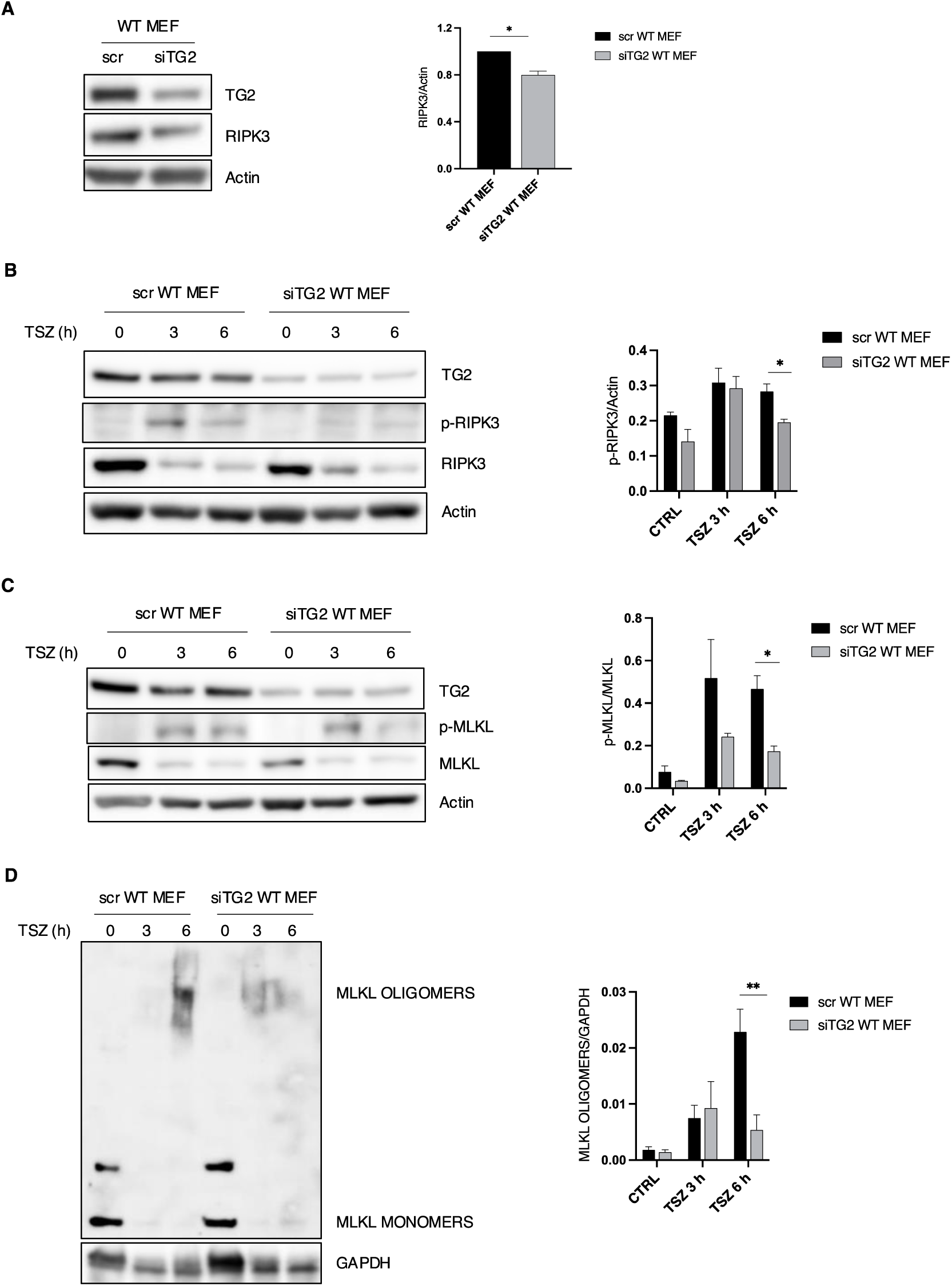
Silencing of TG2 in WT MEFs mitigates necroptosis. (A) Representative western blot and densitometric analysis of RIPK3 in scr WT MEFs and siTG2 WT MEFs in basal conditions. Actin was used as loading control. The graph shows the mean ± SEM of densitometric analysis from three independent experiments. The P-value was determined by unpaired Student’s *t* test (*p < 0.05). (B) Representative western blot and densitometric analysis of p-RIPK3 and TG2 in scr WT MEFs and siTG2 WT MEFs after TSZ treatment for 3 and 6 h. Actin was used as loading control. The graph shows the mean ± SEM of densitometric analysis from three independent experiments. The P-value was determined by unpaired Student’s *t* test (*p < 0.05). (C) Representative western blot and densitometric analysis of p-MLKL, MKLK and TG2 in scr WT MEFs and in siTG2 WT MEFs after TSZ stimulation for 3 and 6 h. Actin was used as loading control. The graph shows the mean ± SEM of densitometric analysis from three independent experiments. The P-value was determined by unpaired Student’s *t* test (*p < 0.05). (D) Representative western blot and densitometric analysis of MLKL oligomers in scr WT MEFs and siTG2 WT MEFs after TSZ treatment for 3 and 6 h to induce necroptosis, the protein samples were treated in a non-reducing condition. GAPDH used as loading control. The graph shows the mean ± SEM of densitometric analysis from three independent experiments. The P-value was determined by unpaired Student’s *t* test (**p < 0.01).

### Effect of TG2 ablation on TNFα signalling

Considering the necroptosis defect in TG2 KO MEF, after TSZ treatment, we wondered whether the lack of TG2 could also affect the upstream regulation of cell death pathways. To this aim, we treated WT and TG2 KO MEFs with TNF to analyze the physiological response to TNFR receptor stimulation. After TNF treatment, we observed the activation of caspase 3 and the cleavage of PARP in cells lacking TG2 (Figure S2A), suggesting defective induction of pro-survival signalling and increased sensitivity to apoptosis. We also analyzed RIPK1, as the main regulator of cell death induction in TNF signalling. Interestingly, RIPK1 interacts with TG2 and of note we detected a cleaved form just in TG2 KO MEFs after TNF (Figure S2B-C). In this regard, it is established that RIPK1 interacts with FADD and caspase-8 to form complex II, which triggers RIPK1-dependent apoptosis. In this context, the cleavage of RIPK1, by caspase-8, is a mechanism for preventing abnormal cell death and necessary for dismantling death-inducing complexes, thereby terminating the death signal [20, 21]. These results suggest that the absence of TG2 not only affects RIPK3 expression and necroptosis but also alters RIPK1 dependent response to TNF.

### H3K4me3 and H3K4me3Q5ser modifications are associated with the *Ripk3* promoter

To define the defective expression of RIPK3 in cells lacking TG2, we studied its epigenetic regulation since its expression can be regulated by DNA methylation [22]. Indeed, it has been demonstrated that the promoter of *Ripk3* is associated to H3K4me3, an histone modification correlated with transcriptionally active chromatin [6, 7]. Interestingly, Farrelly’s group demonstrated that TG2 can covalently bind serotonin (also known as 5-hydroxytryptamine; 5-HT), an excitatory neurotransmitter, to the glutamine 5 of H3K4me3 histone. Specifically, the presence of serotonylation alongside H3K4me3 (H3K4me3Q5ser) enhances the activity of chromatin reader and eraser proteins, stabilizing it and thus favoring gene expression [18].

Considering these premises, we wanted to evaluate whether this histone post-translational modification was present in our model. First, we verified whether TG2 could interact with both H3K4me3 and H3K4me3Q5ser by performing an immunoprecipitation of the histones in the nucleus of WT MEFs. Interestingly, we observed that TG2 interacted with both (Figure 3A-B), thus suggesting that it could trigger the serotonylation of the H3K4me3 histone.

**Figure 3.**
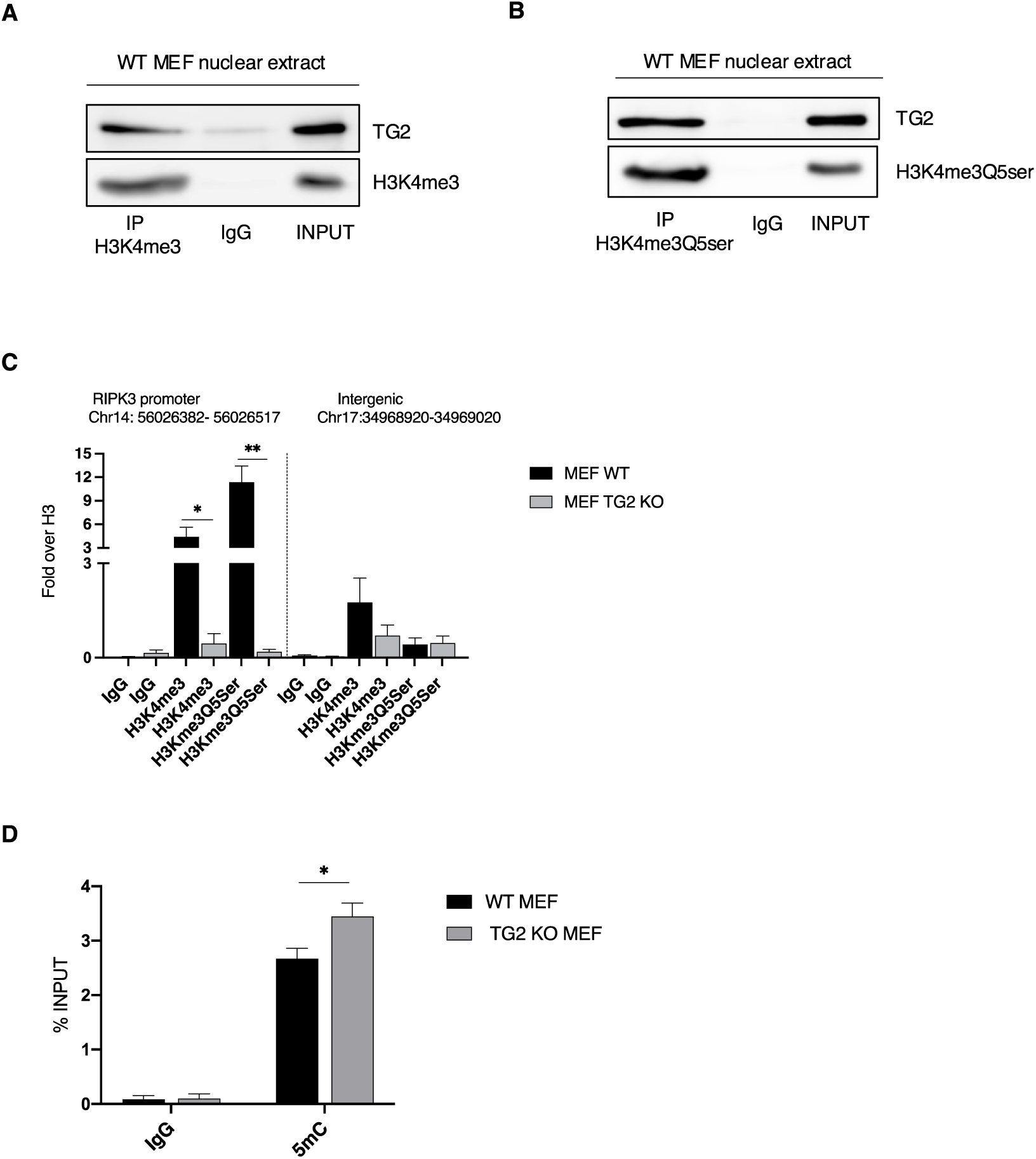
H3K4me3 and H3K4me3Q5ser are associated with the *Ripk3* promoter only in presence of TG2. (A-B) Representative western blot of TG2, H3K4me3 (A) and H3K4me3Q5ser (B) in the nucleus of WT MEFs subjected to immunoprecipitation for H3K4me3 and H3K4me3Q5ser. Cells were lysed, and proteins were immunoprecipitated using anti-H3K4me3 Ab and anti-H3K4me3Q5ser Ab. Input, total cell lysate was used as protein control. (C) ChIP-assay was achieved in WT MEFs and TG2 KO MEFs. ChIP-qPCR analysis was performed in the proximal region of RIPK3 promoter and in an intergenic region, as negative control, using anti-H3K4me3 and H3Kme3Q5Ser antibodies. ChIP signals are normalized as fold over H3. The graph shows the mean ± SEM from three independent experiments. The P-value was determined by two-way ANOVA with Sidak’s multiple comparation test (*p < 0.05, **p < 0.01). (D) Methylation levels of *Ripk3* gene in WT MEFs and TG2 KO MEFs analyzed by MeDIP assay. The graph shows the mean ± SEM from three independent experiments. The P-value was determined by two-way ANOVA with Sidak’s multiple comparation test (*p < 0.05).

Therefore, we wondered whether effectively H3K4me3 and H3K4me3Q5ser histone modifications could be associated with *Ripk3* promoter. To verify this hypothesis, we performed a chromatin immunoprecipitation (ChIP) of both H3K4me3 and H3K4me3Q5ser in WT MEFs and TG2 KO MEFs. Interestingly, we found that H3K4me3 was associated with the promoter region of *Ripk3* gene in WT MEFs, but not in absence of TG2. Of note, we found that H3K4me3Q5ser histone modification was associated to *Ripk3* promoter only in WT MEFs (Figure 3C). In this regard, it is known that serotonylation stabilizes the H3K4me3 mark by inhibiting the activity of histone demethylases [23]. Therefore, in absence of TG2, the lack of serotonylation on H3K4me3 probably favours the removal of methylation on the histone, thus explaining why we did not observe H3K4me3 in TG2 KO MEFs. All together these results indicate that TG2 is essential to promote the serotonylation of H3K4me3 allowing *Ripk3* gene expression, highlighting the potential significance of serotonylation as a contributor to epigenetic modifications [24].

It is known that post-translation modifications of histones also influence DNA methylation. H3K4me3 modification appears to be mutually exclusive with DNA methylation since it prevents the methylation on cytosines in the CpG islands [25–26]. Given this evidence, we wondered whether the absence of the H3K4me3 modification, in cells lacking TG2, was also correlated to *Ripk3* gene methylation. To this aim, we performed the methylated DNA immunoprecipitation (MeDIP) to evaluate the methylation state of *Ripk3* gene and we effectively found it was increased in TG2 KO MEFs, confirming that *Ripk3* promoter is methylated (Figure 3D).

It is also important to mention that serotonylation protects the H3K4me3 from the activity of the histone demethylases JARID1/KDM5 [23] and in turn H3K4me3 histone does not allow DNA methyltransferase to methylate the promoters of the genes associated with it, effectively promoting their expression [23, 24]. In this regard, a proteomic analysis of TG2 nuclear interactors [14] revealed that the enzyme interacts with either demethylase of H3K4me3 (KDM1 and NO66) or methyltransferase (SET1A), further confirming its epigenetic role (Figure S3A).

### 5-Aza-2’-Deoxicytidine (AZA) restores Ripk3 gene expression and necroptosis in cells lacking TG2

To corroborate these results, we verified if the re-expression of RIPK3 in TG2 knock out cells could restore necroptosis pathway. To this aim, we used 5-Aza-2’-Deoxicytidine (AZA), an analogue of cytidine and inhibitor of DNA methyltransferase 1 (DNMT1), able to prevent DNA methylation, allowing gene expression [22]. Thus, we treated TG2 KO MEFs with different concentration of AZA for 4 days to inhibit methylation. Interestingly, we started to observe the rescue of RIPK3 protein expression at 10 μM of AZA (Figure S3B), but the maximum expression of RIPK3 was detected with 20 μM (Figure S3C). These data indicate that *Ripk3* gene is hypermethylated in MEFs lacking TG2 and this is correlated to the absence of H3K4me3 modification.

Finally, we wanted to verify the effect of RIPK3 rescue on necroptosis induction. To this aim, we treated TG2 KO MEFs with AZA for 4 days to restore RIPK3 protein expression levels and then we induced necroptosis by using TSZ for 1, 3 and 6 h. Interestingly, the treatment with AZA and TSZ leads to the rescue of necroptosis process in TG2 KO MEFs as highlighted by the phosphorylation of RIPK3 and MLKL (Figure 4A), the formation of MLKL oligomers (Figure 4B) and the reduced cell viability detected by MTT assay (Figure 4C). These results confirm that the TG2 dependent modification of H3K4me3, prevents *Ripk3* gene methylation, thus favouring its expression and necroptosis induction.

**Figure 4.**
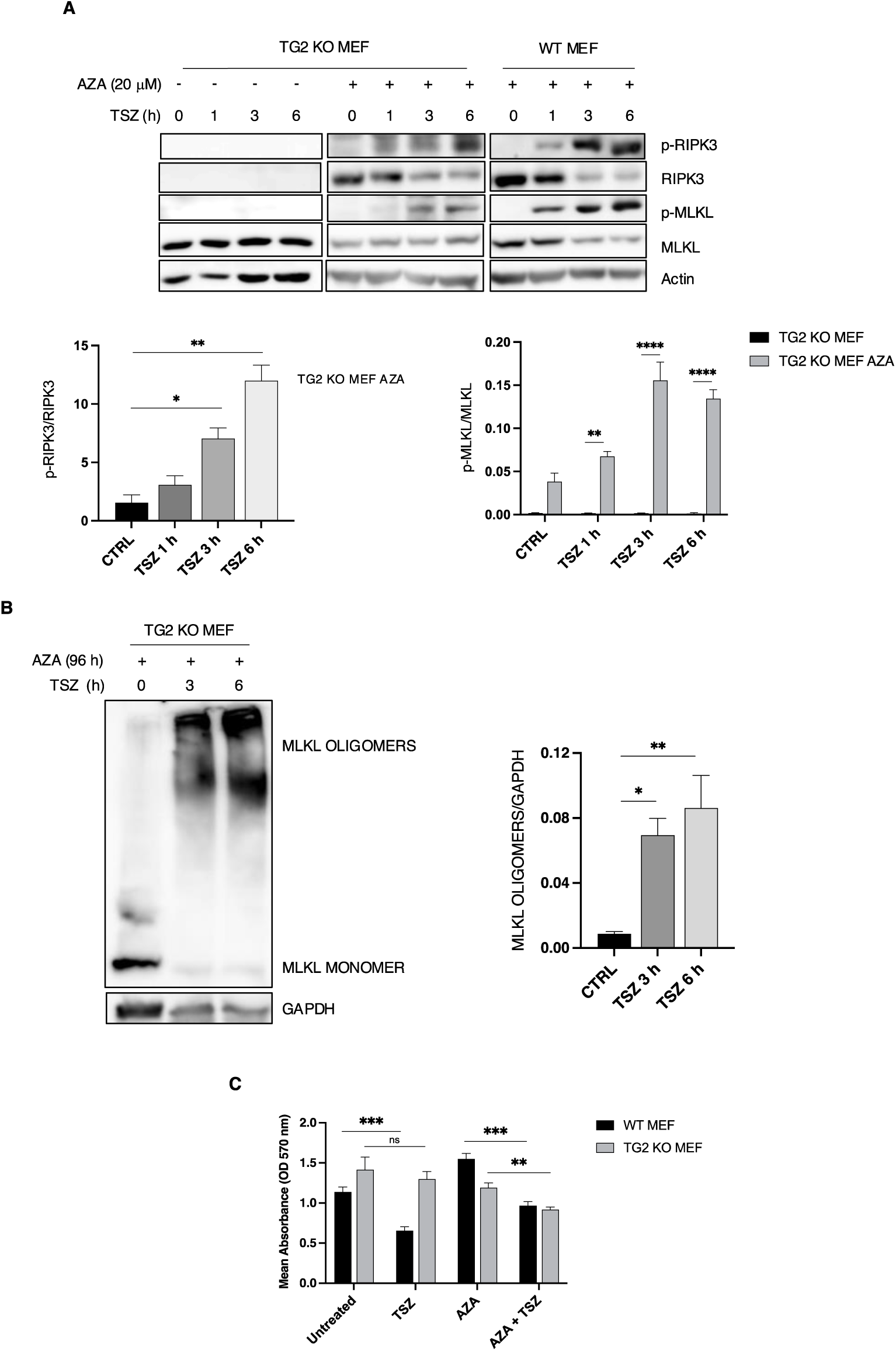
Rescue of necroptosis in TG2 KO MEF treated with AZA. (A) Representative western blot and densitometric analysis of P-RIPK3, RIPK3, P-MLKL and MLKL in WT MEFs and TG2 KO MEFs treated with AZA for 4 days and with TSZ for 1, 3 and 6 h. Actin was used as loading control. The graph shows the mean ± SEM of densitometric analysis from three independent experiments. The P-value was determined by unpaired Student’s *t* test and two-way ANOVA with Sidak’s multiple comparation test (*p < 0.05, **p < 0.01, ****p < 0.0001). (B) Representative western blot analysis and densitometric analysis of the MLKL oligomers and GAPDH in TG2 KO MEFs treated with AZA for 4 days and with TSZ for 3 and 6 h. GAPDH was used as loading control. The graph shows the mean ± SEM of densitometric analysis from three independent experiments. The P-value was determined by unpaired Student’s *t* test (*p < 0.05, **p < 0.01). (C) The graph shows the mean ± SEM of the absorbance (OD 570 nm) of MTT assay from three independent experiments performed on WT and TG2 KO MEFs untreated and treated with AZA 20 μM for 4 days and TSZ for 4 h. The P-value was determined by unpaired Student’s *t* test (**p < 0.01, ***p< 0.001).

In conclusion, our data indicate that TG2 can regulate necroptosis by promoting the expression of *Ripk3,* through a novel epigenetic molecular mechanism. This finding opens new research scenarios that link the enzyme to necroptosis both in its physiological and pathological contexts. Indeed, several studies of the last two decades demonstrated that non-physiological regulation of proteins involved in necroptotic pathway such as RIPK3 can lead to the development of several pathologies such as neurodegenerative disorders, autoimmune diseases, cardiovascular disease, infectious diseases, tumors [27–28]. In this regard, TG2 deregulated expression has been shown to play a key role in cancer progression or inflammation [29–31]. Therefore, it would be of great interest to investigate TG2-dependent modulation of *Ripk3* even in pathological contexts with the aim to use TG2 as a possible target to block RIPK3 improving patient prognosis.

## Acknowledgements

This work was supported in part by grants from the AIRC (21880 to M.P., 24767 to V.P., 27116 to F.R. and 23416 to C.S.), Fondazione Fibrosi Cistica (FFC#8/2022 to M.P.) and the European Union NextGenerationEU through the Italian Ministry of University and Research under PNRR-MAC2-II.3 project PE6 “Heal Italia” CUP E83C22004670001. The Italian Ministry of University and Research (PRIN 2022, CUP E53D23007150006 to F.R.). C.P. was supported by post-doctoral fellowship from Associazione Italiana Ricerca sul Cancro [Project code 28286].

## Author contributions

A.V. and F.R. designed and performed most of the experiments. F.C. performed the immunofluorescence analysis. M.D., L.O. and V.B. helped with the experiments. C.P. and V.P. planned and performed the CHIP assay. F.C. performed the MeDIP assay. F.D. and C.S. contributed to the project. F.R. wrote the paper and together with M.P. conceived the project. All authors read and edited the manuscript.

## METHODS

### Cells and treatments

WT MEFs and TG2 KO MEFs were obtained by spontaneous immortalization of fibroblasts derived from C57BL/6 mice embryos at E14 either wild type or knockout for TG2 [32]. Cells were cultured in Dulbecco’s modified Eagle’s medium (Lonza) supplemented with 10% fetal bovine serum, 100 μg/ml streptomycin and 100 units/ml penicillin, at 37°C and 5% CO2 in a humidified atmosphere. To induce necroptosis, MEFs were pre-treated with 20 mM Z-VAD-FMK (Invivogen) and 2 mM Smac mimetic (Sigma-Aldrich) also known as antagonists of IAPs for 1 h. After 1 h 150 ng/ml tumour necrosis factor a (TNFa) (BioLegend) was added to the media for 1, 3, 4 and 6 h. WT MEFs were silenced for TG2 by Small Interference RNA (siRNA) using Lipofectamine 2000 (Invitrogen) according to the manufacturer’s instructions. The siRNA for TG2 and scramble sequences were obtained from OriGene company (SR419141). To induce *Ripk3* gene expression in TG2 KO MEFs, a concentration of 10 and 20 mM 5-Aza-2’-Deoxicytidine (AZA) (Sigma-Aldrich) an inhibitor of DNA methyltransferase 1 (DNMT1) was used for 4 days.

### Western blot

MEF cells were washed with ice-cold PBS and then collected in lysis buffer consisting of 20 mM Tris–HCl at pH 7.4, 150 mM NaCl, and 1% Triton X-100 supplemented with a cocktail of protease inhibitors. The protein concentrations were quantified using the Bradford assay with bovine serum albumin as a reference standard. Portions of the total protein extracts, obtained from cells subjected to various treatments, were denatured. To analyse the oligomeric form of MLKL, protein extracts were processed in a non-reducing condition. Aliquots of protein extracts were separated by SDS– polyacrylamide gel electrophoresis and subsequently transferred onto a nitrocellulose membrane. After blocking with 5% non-fat dry milk in T-PBS (PBS containing 0.05% Tween-20) for 1 h at room temperature, the membranes were then incubated overnight with the specified primary antibodies. Following this, the membranes were exposed to an HRP-conjugated secondary antibody for 1 h at room temperature, and the resulting signal was visualized using Immobilon Western detection reagents from Millipore.

### Immunoprecipitation

Nuclear fraction from MEF cells was obtained using the NE-PER Nuclear and Cytoplasmic Extraction Kit from (Thermo Scientific). Nuclei were lysed in a buffer containing 150 mM NaCl, 50 mM Tris–HCl pH 7.5, 2 mM EDTA, 2% NP-40 and freshly added protease inhibitor cocktail. An amount of 500 μg of nuclear proteins was subjected to immunoprecipitation using 4 μg of specific antibodies in combination with 15 μl of DynabeadsTM Protein G (Invitrogen), according to the manufacturer’s instructions. LDS Sample Buffer 2× (Life Technologies) containing 2.86 M 2- mercaptoethanol (Sigma-Aldrich) was added to beads, and samples were boiled at 95°C for 10 min. Supernatants were analysed by Western blot.

### Chromatin immunoprecipitation

WT MEFs and TG2 KO MEFs were cross-linked in 1% (vol/vol) formaldehyde to the culture medium for 10 min at room temperature and then quenched with 125 mM glycine for 5 min. Cross-linked cells were washed twice with PBS, scraped, and pelleted by centrifugation at 1300 x g for 10 min. Cell pellet was lysed in nuclei extraction buffer (5 mM Pipes pH 8, 85 mM KCl, 0.5% NP40) for 2 h at 4°C under rotation. The lysate was centrifugated at 1200 x g for 5 min at 4°C. Nuclei pellet was resuspended in sonication buffer (10 mM ethylenediaminetetraacetic acid (EDTA) pH 8, 50 mM Tris–HCl pH 8, SDS 1%) and the chromatin was sheared by sonication with Bioruptor (Diagenode) to obtain chromatin size between 100 and 1000 base pairs. The size of the sheared chromatin was confirmed by 0.8% agarose gel electrophoresis compared to unsheared chromatin. Share DNA was then quantified, diluted 1:10 with dilution buffer (0.01% SDS, 1.1% Triton X100, 1.2 mM EDTA, 16.7 mM Tris/HCl pH 8.0, 167 mM NaCl) and 40 μg of chromatin was immunoprecipitated with 2 μg of anti-H3K4me3, anti-H3K4me3Q5ser or anti-H3 primary antibodies overnight at 4°C by head-to-head rotation. IgGs (Sigma-Aldrich) were used as a negative control. Dynabeads protein G (Invitrogen, Life technologies) were incubated with the mixture under rotation at 4°C for 2 h, then washed three times with low salt buffer (0.1% SDS, 2 mM EDTA, 1% Triton, 20 mM Tris pH 8, 150 mM NaCl), high salt buffer (0.1% SDS, 2 mM EDTA, 1% Triton, 20 mM Tris pH 8, 500 mM NaCl) and Tris-EDTA buffer (1 mM EDTA, 10 mM Tris pH 8) for 5 min each. Precipitated material was then treated with RNase inhibitors cocktail (AM2288; Ambion) for 1 h at 37°C to remove RNA contamination and heated at 65°C overnight to reverse formaldehyde cross-links. De-crosslinked chromatin was than treated 0.6 mg/ml proteinase K (Sigma Aldrich) at 55°C for 2 h. Eluted DNA was recovered by Phenol-Chloroform mediated purification and analyzed by qPCR analysis with primers directed against the *Ripk3* promoter region (mRipk3 sense 5′-AACTGCTTCCACCCGAGAAG-3′, mRipk3 antisense 5′-AAAATGTGGTTGGCAAGCGG-3′). A primer set amplifying an *HSP70.3* intergenic region (mHSP70.3 Intergenic Region sense 5′- GTGGCGCATGCCTTTGAT-3′; mHSP70.3 Intergenic Region antisense 5′- CTTTGTAGAACAGGCTGACCTTGA-3′) was used as negative control.

### Methylated DNA Immunoprecipitation (MeDIP)

Genomic DNA (8 ug) isolated with Quick-DNA Miniprep Kit (Zymo Research) were diluted in 400 µl of water and sonicated (40% amplitude; 0.5 cycle) to obtain fragments about 500bp-300bp. DNA samples were denatured for 10 min at 95°C and then cooled on ice for 10 min. From 3 to 10% of the volume DNA was stored as Input DNA while the remaining part was diluted in IP buffer 2X (20 mM Na-Phosphate buffer pH 7.0, 0.28 M NaCl, 0.1% Triton X-100) and divided into two vials containing 4 μg of anti-5mC (EpiGentek) or normal IgGs (Santa Cruz Biotechnology) as controls. DNA– antibody mixtures were incubated overnight on a rotating platform and then 45 μl of salmon sperm-saturated Protein-A/G Agarose beads (Millipore) were added and incubated for 2 h at 4°C. Beads and immunocomplexes were washed three times with 1X IP buffer at 4°C on a rotating platform for 5 min, incubated in 250 μl of digestion buffer (50 mM Tris-HCl pH 8, 10 mM EDTA, 0.5% SDS) containing 50 μg proteinase K (Sigma-Aldrich) for 2 h at 55°C and subjected to standard DNA precipitation procedure with phenol–chloroform-isoamyl alcohol solution in the presence of 15 μg of glycogen during the ethanol precipitation. Air-dried DNA pellets were resuspended in 30 μl water and 4 μl were used for qPCR amplification. The percent input method was used for the analysis of results according to 100*2^ (Adjusted input - Ct (IP) where Adjusted input is the Raw Ct Input - log2 of the Input dilution factor. Primers used for the analysis were: mRipk3 sense 5′- AACTGCTTCCACCCGAGAAG-3′, mRipk3 antisense 5′-AAAATGTGGTTGGCAAGCGG-3′.

### Quantitative RT-PCR

RNA from MEFs was extracted using Universal RNA Purification Kit (Cat# E3598 EUR_X_) according to the manufacturer’s instructions. 1 μg of RNA was reverse transcribed to cDNA using SensiFAST^TM^ cDNA Synthesis Kit and used for quantitative real-time PCR (RT-qPCR) experiment using SensiFAST SYBR Hi-ROX Kit following manufacturer’s instructions. Thermocycling consisted of initial polymerase activation at 98°C for 5 min, followed by 35 cycles of 95°C for 15 sec, 68°C for 10 sec, and 72°C for 20 sec. Data acquisition was performed at this stage, and the reaction was finished by the built-in melt curve. Relative amounts of mRNA were calculated using the comparative Ct method. The following primers were used in this study. mActin sense: 5′- GGCTGTATTCCCCTCCATCG - 3′, mActin antisense 5′- CCAGTTGGTAACAATGCCATGT -3′; mRipk3 sense: 5′- GAAGACACGGCACTCCTTGGTA -3′, mRipk3 antisense 5′- CTTGAGGCAGTAGTTCTTGGTGG -3′.

### Immunofluorescence analysis

MEFs were fixed in 4% formaldehyde for 10 min at room temperature (RT), washed with PBS, and blocked with 5% bovine serum albumin and 0.1% Triton X-100 (Sigma-Aldrich). The following primary antibodies were used: rabbit polyclonal antibody against MLKL, mouse antibody against Actin. Cells were then incubated with the appropriate secondary antibodies, conjugated with Alexa Fluor 488 or Alexa Fluor 555 (Invitrogen). Nuclei were stained with 1 μg/mL Hoechst 33342 (Invitrogen), and slides were observed at microscope. Images were captured by Zen Pro Microscopy software, and representative images were composed in an Adobe Photoshop CS6 format (Adobe Systems Inc., San Jose, CA, USA).

### MTT assay

To measure cellular metabolic activity as an indicator of cell viability, the MTT assay (M5655, Merck KGaA, Germany) was performed. 50000 cells/well were plated in a 96-well plate and incubated at 37°C. After 24 h, 50 µl of MTT (5 mg/ml) was added to each well and cells were incubated at 37°C for 2.5 h. Then presence of formazan crystals by metabolically active cells was checked and 150 µl of DMSO solvent per well was added. The plate was orbitally shaken in the dark for 15 min, then read with absorbance at OD570 nm using Byonoy Absorbance 96 plate reader (Hamburg, Germany). Representative images showing the presence of purple formazan crystals were composed in an Adobe Photoshop CS6 format (Adobe Systems Inc., San Jose, CA, USA).

### Antibodies

Anti-Actin (A2066) Sigma-Aldrich; anti-GAPDH (G9545) Sigma-Aldrich; anti-Histone H3K4me3 (AB8580) abcam; anti-Histone H3K4me3Q5ser (ABE2580) Merck Millipore; anti-Histone H3 (17168-1-AP) proteintech; anti-MLKL (D6W1K) Cat#37705 cell signalling; anti-phospho-MLKL (D6E3G) Cat#37333 cell signaling; anti-phospo-RIPK1 (E7G60) Cat#53286 cell signaling; anti-RIPK3 (NBP1-77299) Novus Biologicals; anti-phospo-RIPK3 (T231/S232) Cat#91702 cell signaling; anti-TGM2 (D11A6) Cat#3557 cell signaling.

### Statistical analysis

GraphPad was used for statistical analysis. ImageJ64 software was used for densitometric analysis. Statistical significance was determined using the Student’s t-test or one-way ANOVA test. P-values smaller than 0.05 (P < 0.05) were considered to be significant.

---

**Figure S1.**
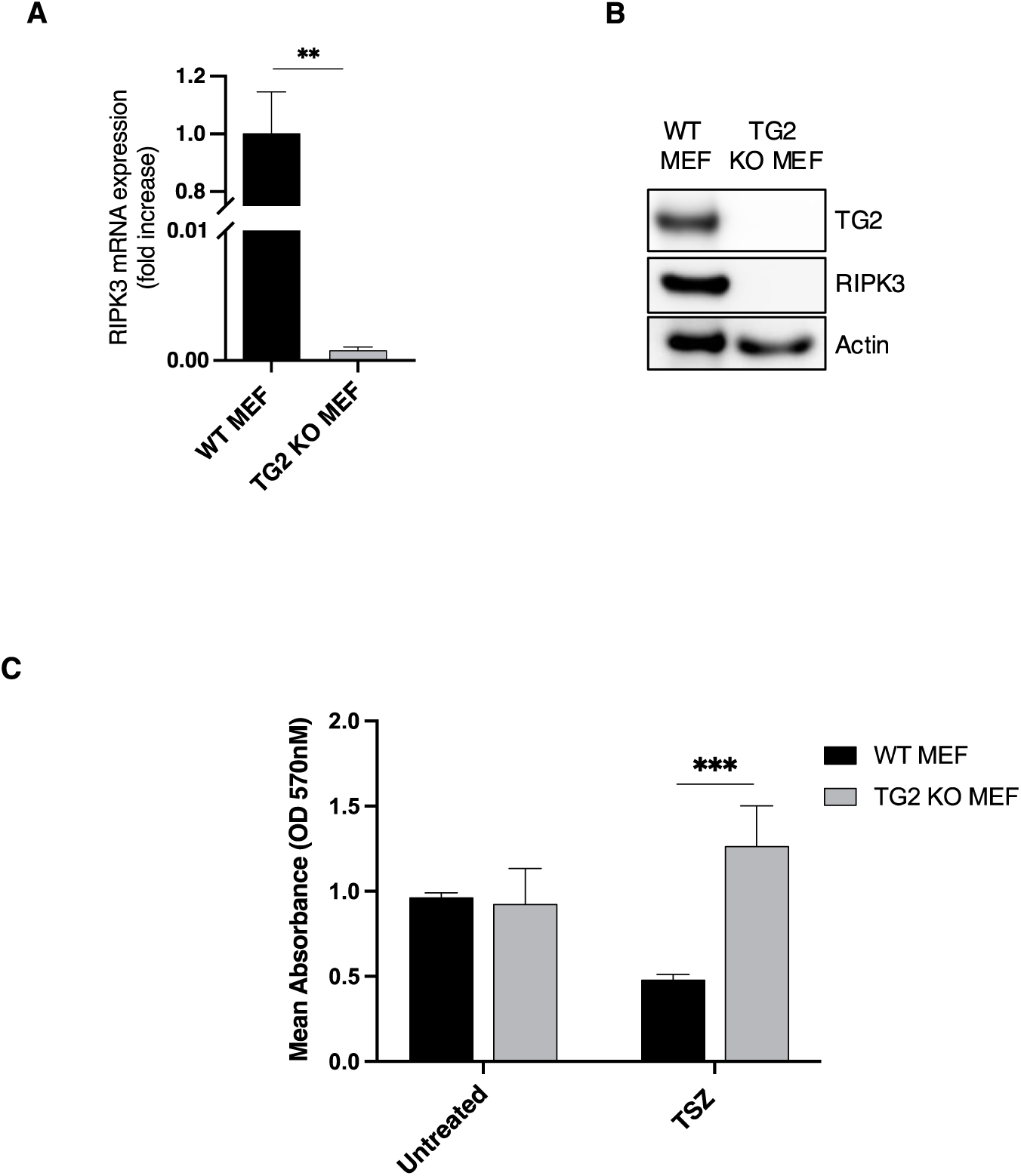
(A) RIPK3 mRNA levels, quantified by RT-qPCR in WT and TG2 KO MEFs in basal conditions normalized on actin levels. The graph shows the mean ± SEM of the mRNA levels from three independent experiments. The P-value was determined by unpaired Student’s *t* test (**p < 0.01). (B) Representative western blot analysis of RIPK3 in WT and TG2 KO MEFs in basal conditions. Actin was used as loading control. (C) The graph shows the mean ± SEM of the absorbance (OD 570 nm) of MTT assay from three independent experiments performed on WT and TG2 KO MEFs untreated and treated with TSZ for 4 h. The P-value was determined by two-way ANOVA with Sidak’s multiple comparation test (***p< 0.001).

**Figure S2.**
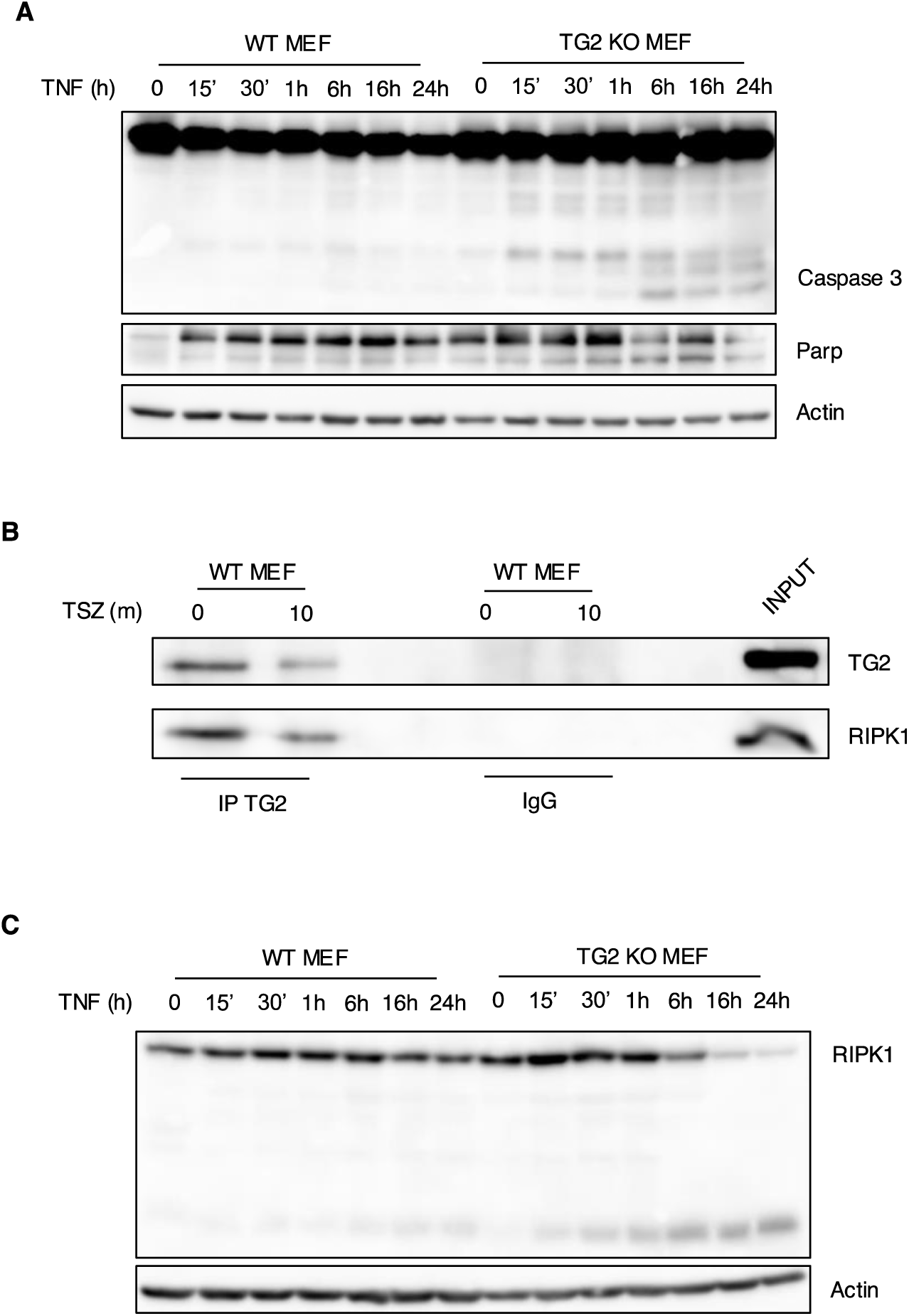
(A) Representative western blot analysis of caspase 3 activation and PARP cleavage in WT and TG2 KO MEFs after TNF treatment. Actin was used as loading control. (B) Representative western blot of TG2 and RIPK1 in WT MEFs subjected to immunoprecipitation for TG2. Cells were lysed, and proteins were immunoprecipitated using anti-TG2 Ab. Input, total cell lysate was used as protein control. (C) Representative western blot analysis of RIPK1 in WT and TG2 KO MEFs after TNF treatment. Actin was used as loading control.

**Figure S3.**
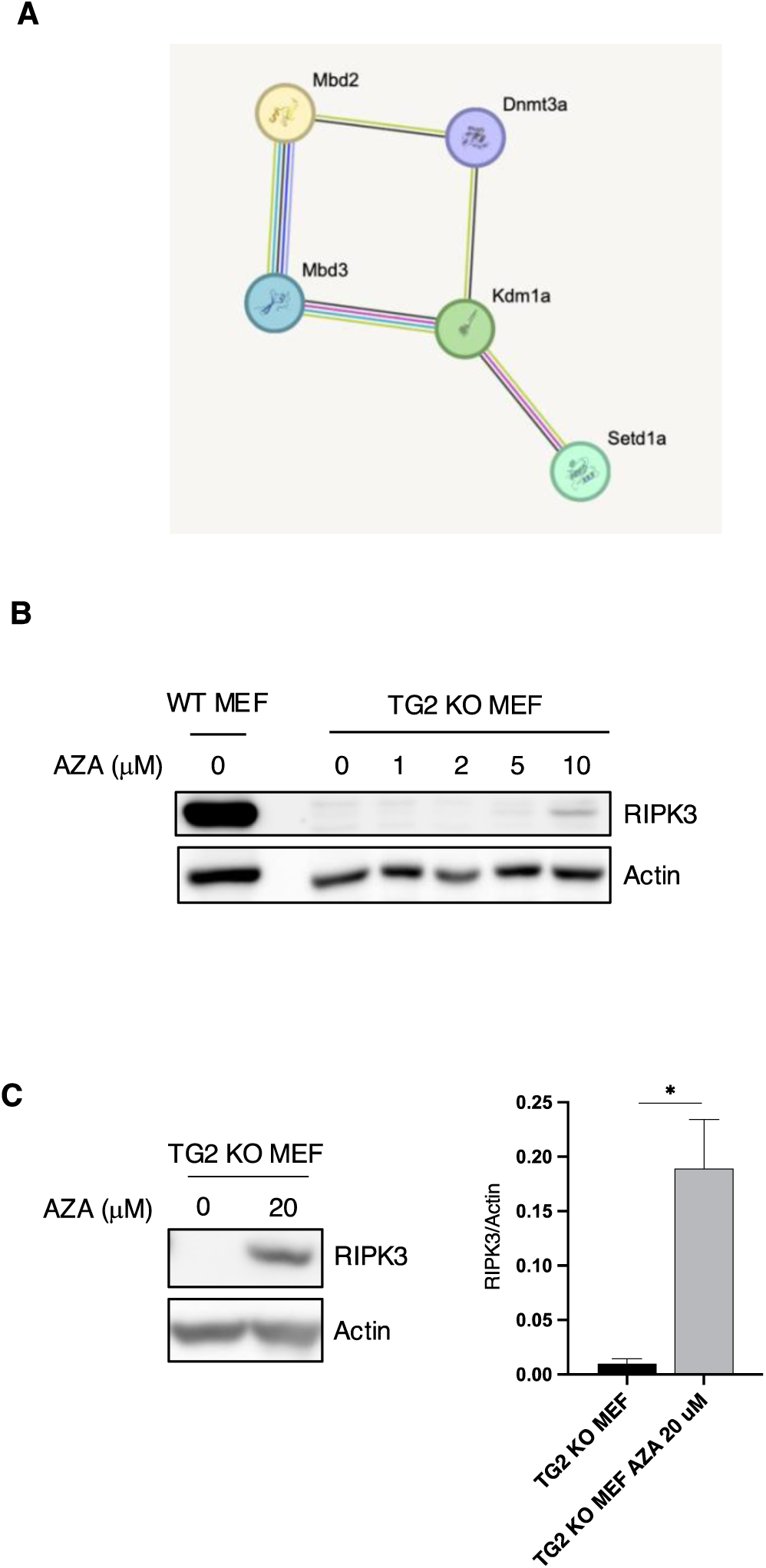
(A) Protein-protein interaction network of the DNA methylases and demethylases identified in the TG2 nuclear interactome analysis. The network, containing identified proteins, was mapped using the STRING system (http://string-db.org/). (B) Representative western blot of RIPK3 in WT MEFs and TG2 KO MEFs treated with different concentrations of AZA for 4 days. Actin was used as loading control. (C) Representative western blot and densitometric analysis of RIPK3 in TG2 KO MEFs treated with AZA 20 μM for 4 days. Actin was used as loading control. The graph shows the mean ± SEM of densitometric analysis from three independent experiments. The P-value was determined by unpaired Student’s *t* test (*p < 0.05).

